# Engineered Receptors for Soluble Cell-to-Cell Communication

**DOI:** 10.1101/2024.09.17.613377

**Authors:** Dan I. Piraner, Mohamad H. Abedi, Maria J. Duran Gonzalez, Adam Chazin-Gray, Iowis Zhu, Pavithran T. Ravindran, Thomas Schlichthaerle, Buwei Huang, David Lee, David Baker, Kole T. Roybal

## Abstract

Despite recent advances in mammalian synthetic biology, there remains a lack of modular synthetic receptors that can robustly respond to soluble ligands and in turn activate cellular functions. Such receptors would have extensive clinical potential to regulate the activity of engineered therapeutic cells, but to date only receptors against cell surface targets have approached clinical translation. To address this gap, we developed a receptor architecture based on synNotch, called SyNthetic Intramembrane Proteolysis Receptor (SNIPR), that has the added ability to be activated by soluble ligands, both natural and synthetic, with remarkably low baseline activity and high fold activation. SNIPRs are able to access an endocytic, pH-dependent cleavage mechanism to achieve soluble ligand sensing, in addition to employing a canonical-like pathway for detecting surface-bound ligands. We demonstrate the therapeutic capabilities of the receptor platform by localizing the activity of CAR T-cells to solid tumors where soluble disease-associated factors are expressed, bypassing the major hurdle of on-target off-tumor toxicity in bystander organs. We further applied the SNIPR platform to engineer fully synthetic signaling networks between cells orthogonal to natural signaling pathways, expanding the scope of synthetic biology. Our design framework enables cellular communication and environmental interactions, extending the capabilities of synthetic cellular networking in clinical and research contexts.

---

The fundamental basis of biochemical signal transduction lies in the ability of cells to produce, sense, and react to small diffusible molecules. This capability allows them to coordinate intricate functions and respond to environmental stimuli beyond their immediate vicinity. For instance, morphogens play a critical role in shaping the three-dimensional pattern of an embryo during development^1^, while cytokines are involved in shaping cell state changes and recruiting a diverse array of immune cells to the site of disease^2^. The ability to mimic these natural systems and interface with soluble factors in synthetic biological systems would enable engineered cells to integrate signals from distant sources and activate therapeutic programs. Furthermore, artificial signaling molecules could provide privileged communication channels between cells, a means to specifically engage engineered cells after administration to patients. Using these bioorthogonal channels of communication, engineered cells could promote large-scale cellular coordination, facilitating complex multicellular behavior and delivering synergistic therapeutic benefits.

Despite this potential, progress toward developing sensitive and robust biosensors for soluble factors has been limited. The MESA receptor platform represents an innovation in this field^3^, but achieving high sensitivity and dynamic range in therapeutic cell types remains a challenge. Furthermore, MESA relies on viral components and a multi-chain architecture that make it a difficult platform to translate. TanGo^4^ and ChaCha^5^ are GPCR-fused protease architectures that offer an alternative approach but still require multiple components, which can constrain the use of therapeutic delivery vehicles like lentiviral or adeno-associated viral vectors. Moreover, these GPCR-based designs lack the flexibility in ligand selection that is inherent to receptors using modular binding domains (e.g., scFvs). The OCAR platform also relies on co-delivery of two receptor chains to allow for ligand-induced dimerization and activation^6^. A compact, single-chain receptor capable of modularly sensing soluble factors would overcome the limitations of current systems and unlock the potential for engineered receptors to coordinate therapeutic genetic programs in clinically-relevant cell types.

Ideally, receptors used for soluble factor detection must operate with high fidelity, generating strong signals in the ON state while minimizing basal signaling in the OFF state. These receptors should also be compact to enable efficient delivery to relevant immune cell types and minimize potential issues related to subunit stoichiometry. The synthetic Notch Receptor (synNotch) represents a prototypical engineered receptor, employing cleavage by endogenous γ-secretase to release its transcription factor upon binding a cell surface ligand and demonstrating robust activation in therapeutically relevant cell types including Chimeric Antigen Receptor (CAR) T-cells^7,8^. Recently, a new receptor with a Notch-based architecture, the SyNthetic Intramembrane Proteolysis Receptor (SNIPR), was developed. SNIPRs provide a compact and readily tunable scaffold for custom signal transduction along the Notch cleavage paradigm^9^. The ability of SNIPRs to robustly and selectively respond to ligand binding despite the omission of the Notch LNR regulatory domains, widely thought to be the key determinants of Notch activation, implies that SNIPRs may employ an alternate signaling pathway that bypasses the mechanosensing filter which precludes Notch and synNotch from detecting soluble ligands. We set out to investigate whether the SNIPR system could be adapted to sense soluble ligands, which could enable a wide variety of applications (**Figure 1a**).

**Figure 1:**
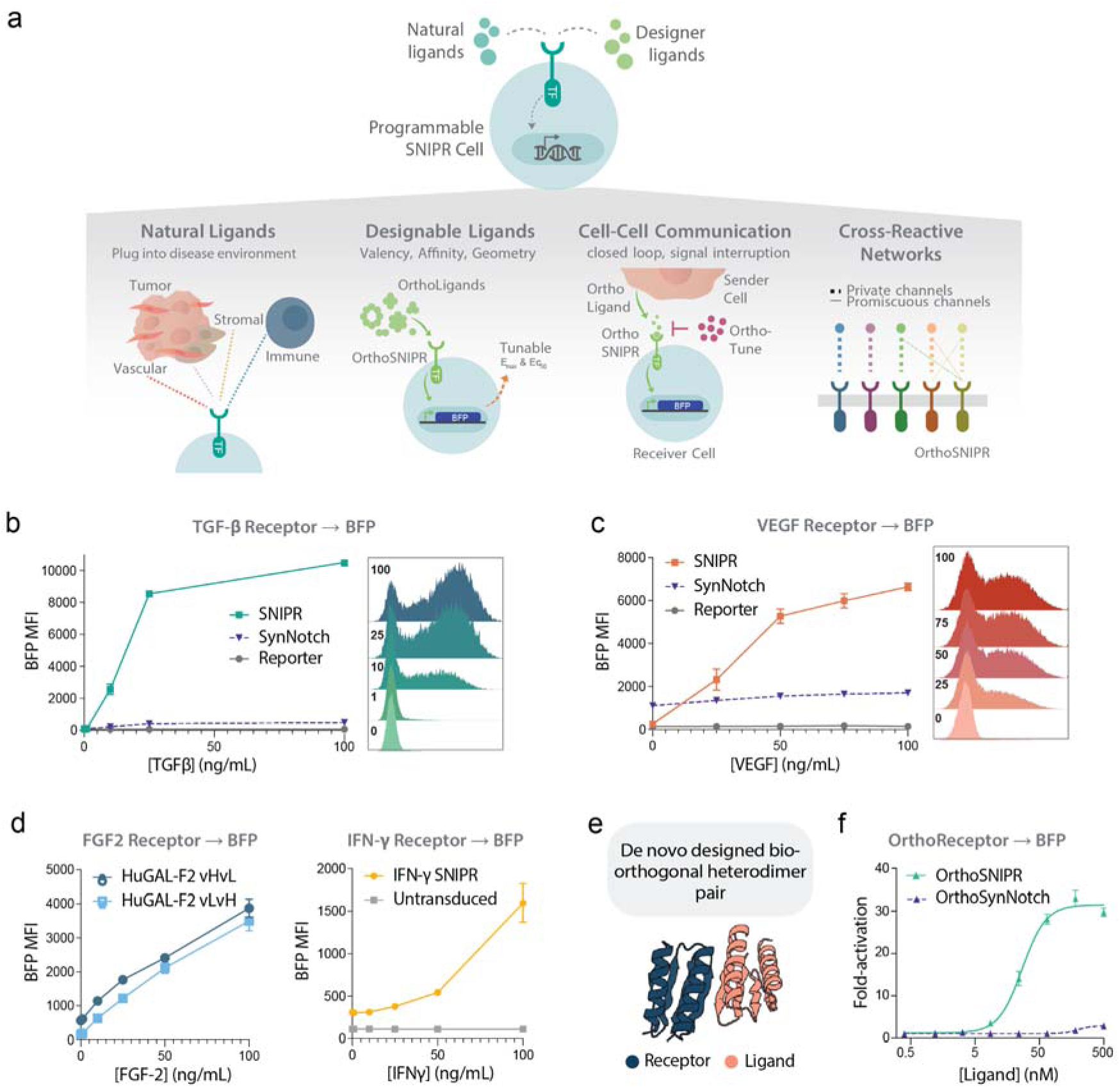
Engineering versatile synthetic receptors capable of sensing natural and engineered soluble ligands. **a)** Clinical and research applications for engineered receptors capable of soluble ligand detection. **b)** Left: Activation of a TGF-β Responsive SNIPR or synNotch driving a BFP reporter circuit in primary human CD3^+^ T-cells upon addition of recombinant human TGF-β1 (N = 2; +/- SD). Right: Representative histograms of reporter expression level at the indicated TGF-β concentrations. **c)** Induction (left) and representative histograms (right) of a VEGF sensing SNIPR-to-BFP circuit in primary human CD3^+^ T-cells via recombinant human VEGF (N = 3; +/- SD). **d)** Receptor-mediated induction of a BFP reporter in primary human CD3^+^ T-cells using recombinant human FGF-2 (left) or IFN-γ (right) (N = 2 and 3, respectively; +/- SD). **e)** A structural model of the *de novo* designed heterodimer used to establish a bio-orthogonal receptor-ligand pair. **f)** Fold activation over background of an orthogonal ligand responsive SNIPR or synNotch driving a BFP reporter circuit in Jurkat T-cells via introduction of ortho-ligand C1-active at concentrations ranging from 0.4 to 500 nM (N = 3; +/- SEM).

**Figure S1:**
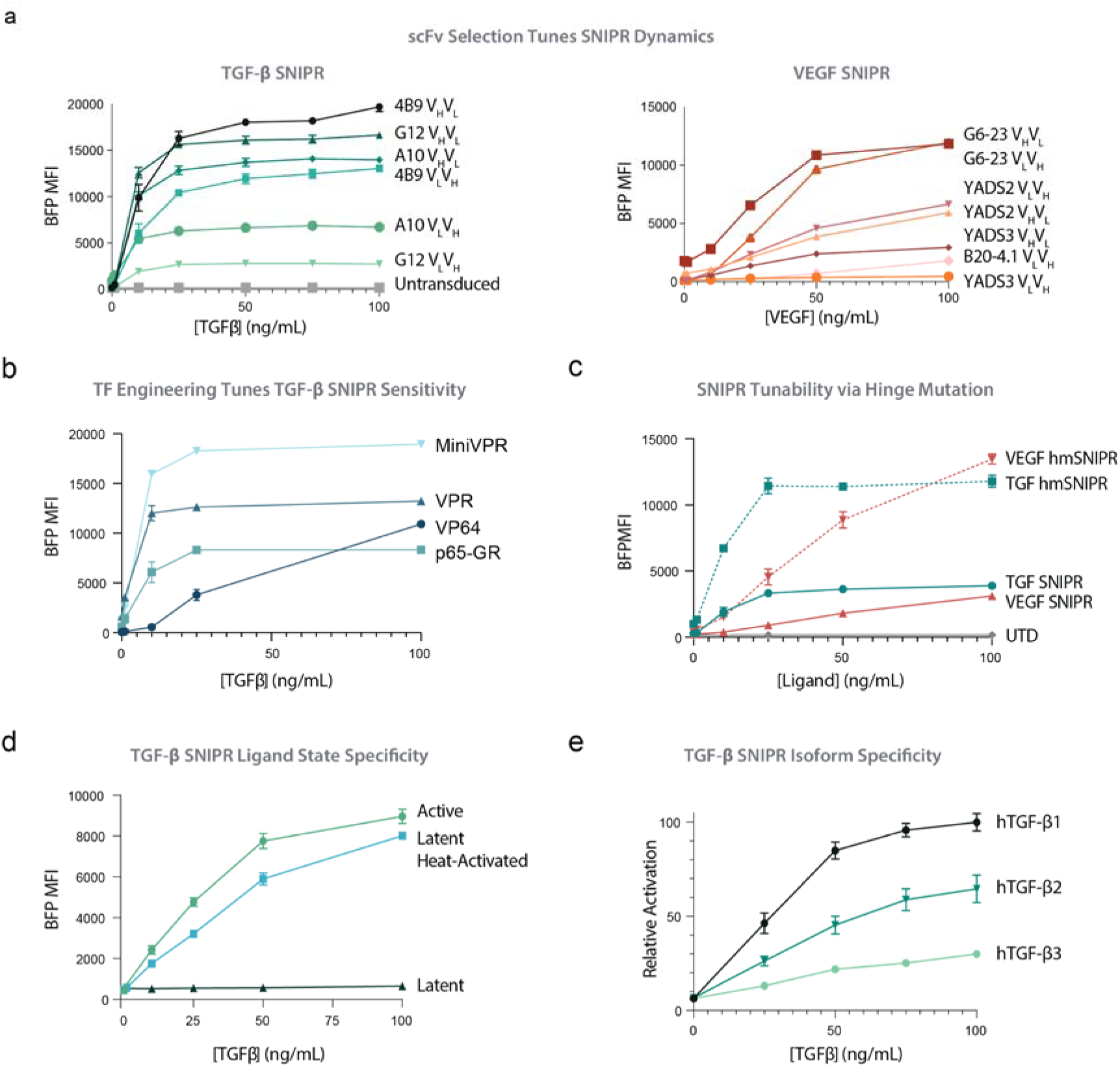
**a)** BFP reporter induction in a panel of SNIPR variants bearing scFvs targeted against TGF-β (left) or VEGF (right) in V_H_V_L_ or V_L_V_H_ orientation. Data describe primary human CD3^+^ T-cells, N = 3 (TGF) or 2 (VEGF); +/- SD). **b)** BFP reporter gene induction of primary human CD3^+^ T-cells expressing SNIPRs containing a set of unique Transcription Activation Domains (TADs) (N = 2; +/- SD). **c)** BFP reporter induction by conventional or hinge-mutant (hm) TGF and VEGF SNIPR or untransduced T-cells (N = 3; +/- SD). **d)** TGF SNIPR activation in primary human CD3^+^ T-cells using recombinant active TGF-β, recombinant latent TGF-β, or recombinant latent TGF-β re-activated via heat exposure (N = 3; +/- SD). **e)** TGF SNIPR activation in primary human CD3^+^ T-cells using recombinantly expressed human TGF-β1, β2, or β3 protein isoforms (N = 3; +/- SD).

Our findings confirm that SNIPRs can utilize an alternative signaling pathway, allowing for robust and selective responses to soluble ligand binding. SNIPRs possess the unique capability to interact with a range of physiological or synthetic ligands, whether they are tethered or soluble. This versatile feature makes them an asset to the current array of cell-engineering tools, with potential applications in cancer therapy, and engineering development. Here we have shown that T cells engineered with SNIPR technology can effectively react to soluble factors and drive therapeutic payload production in solid tumor animal models. Moreover, we also show that SNIPRs can serve as a fundamental building block for constructing modular bio-orthogonal communication systems in cells that can operate independently of natural signaling pathways.

## SNIPRs Enable Sensing of Natural and Engineered Soluble Factors

SNIPRs remain inactive in the absence of ligands but exhibit a robust transcriptional response when membrane-bound ligands are encountered, despite lacking the LNR domains thought to be critical for regulating Notch activation. We hypothesized that SNIPRs may activate through an alternative pathway to the conventional force-induced LNR stretching model^10^, enabling the detection of soluble factors that are inaccessible to Notch family receptors. To assess this potential, we initially designed SNIPRs with extracellular domains (ECDs) incorporating scFvs derived from a set of antibodies against the broadly tumor-associated cytokines Transforming Growth Factor β (TGF-β)^11^ and Vascular Endothelial Growth Factor α (VEGF)^12^ (**Supplementary Figure 1a**). TGF-β is a critical tumor signaling molecule that is intrinsically tumorigenic, suppressive against tumor-infiltrating immune cells, and broadly expressed across a variety of solid cancers^13^. Likewise, VEGF is a pleiotropic tumor-associated signaling molecule that can modulate the immune environment but is best known for its role in stimulating angiogenesis within the tumor^12^. We found that primary human CD3+ T-cells bearing TGF-β SNIPR driving BFP reporter circuits signaled robustly upon application of recombinant activated TGF-β. Similarly, the VEGF SNIPR panel displayed robust signaling upon titration of recombinant VEGF. As expected, synNotch receptors bearing the same anti-TGF-β or anti-VEGF extracellular domains failed to signal upon addition of the cognate ligands (**Figure 1b,c**). We demonstrated the tunability of the SNIPR by varying both the scFv identity and orientation (**Supplementary Figure 1a**) and the potency of the receptor’s transactivation domain (**Supplementary Figure 1b**). We also found that receptor activity can be modulated by replacing the cysteine residue in the hinge domain with a serine residue (**Supplementary Figure 1c**). The TGF-β SNIPR is specific to the active form of TGF-β (**Supplementary Figure 1d**), suggesting its potential for tumor microenvironment detection^14^, and was cross-reactive across human TGF-β isoforms, albeit with a preference for TGF-β1 (**Supplementary Figure 1e**). To expand the range of possible soluble targets in disease environments, we developed functional SNIPRs capable of detecting the stromal signaling factor Fibroblast Growth Factor 2 (FGF-2) as well as Interferon-γ (IFN-γ), a key inflammatory cytokine (**Figure 1d**). Collectively, these findings indicate that SNIPRs can serve as a versatile receptor architecture for detecting cancer-related and inflammatory soluble signaling molecules.

We next investigated whether the SNIPR system could be adapted to enable bio-orthogonal cell signaling in Jurkat T-Cells. We designed SNIPRs that have one subunit of a designed “LHD” heterodimers^15^ as its extracellular domain, and evaluated signaling in response to the soluble heterodimeric partner (**Figure 1e,f**). Addition of the cognate subunit resulted in strong transcriptional response. In contrast, no signaling was observed when the LHD heterodimer was incorporated into the original synNotch system–the Notch LNR domains effectively block signaling through soluble ligands. This “OrthoSNIPR” system is bio-orthogonal since the LHD subunits do not interact with naturally occuring proteins, and hence enables the creation of “private” channels of communication between engineered cells that could improve therapeutic efficacy and safety. We next investigated the tunability of OrthoSNIPR signaling. We generated ligands with the same binding domain but differing valency and geometry, by fusing them to a series of designed oligomeric scaffolds^16^ (**Figure 2a**). These ligands generate a range of signaling strengths as reflected in both EC_50_ and E_max_, with higher valency ligands generally having increased sensitivity, cooperativity, and maximal activation up to a 40-fold increase over the baseline level (**Figure 2b**). The OrthoSNIPR recapitulated the tunability of the original receptor by mutation of the hinge cysteine to a serine residue (**Supplementary Figure 2a**). The ability to tune signaling using a series of ligands with graded receptor activation should be useful in refining engineered signaling networks. Engineered signaling pathways that are conditional on the presence of additional factors could have considerable therapeutic utility. For example, if CAR T cells detect a signal from other engineered cells indicating they are in the wrong environment, or if patients experience CRS symptoms, off- or damper-switches could mitigate toxicity. To enable such conditional signaling, we designed a weakly homomeric ligand that strongly induces receptor signaling unless another engineered factor, orthoTuner, is present in the environment. OrthoTuner has higher affinity for the subunits of the homodimeric ligand than they have for each other, and hence forms heterodimers that contain only one receptor interacting LHD module (**Figure 2c**). Addition of OrthoTuner modulates signaling in a dose-dependent manner (**Figure 2d**). This conditionality enables modulation of the strength of OrthoSNIPR signaling by external inputs or communication between engineered cells.

**Figure 2.**
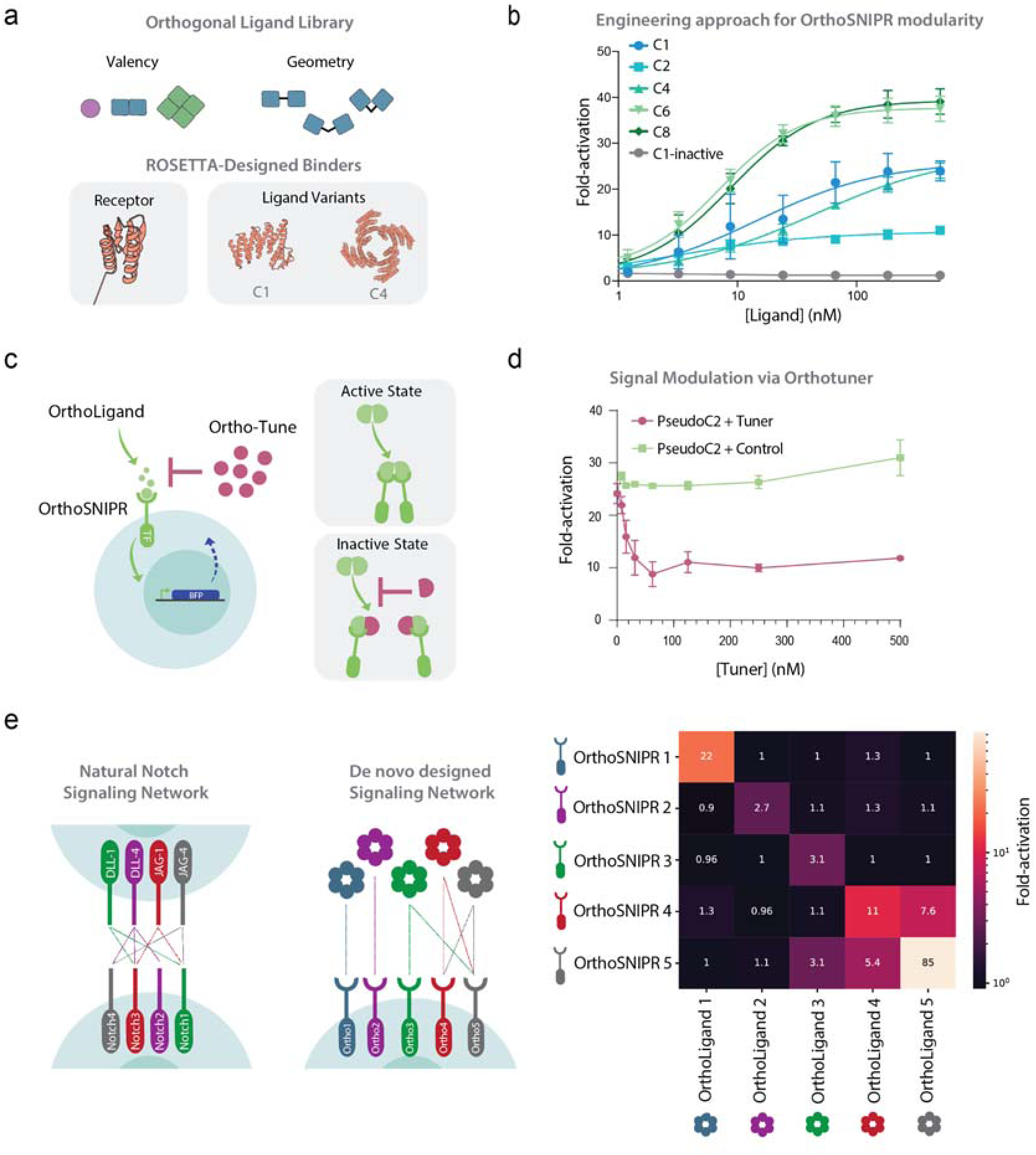
Construction of bio-orthogonal orthoSNIPR signaling systems. **a)** Design of a series of synthetic ligands with a range of geometries and valencies. **b)** Activation of an ortho-ligand responsive SNIPR driving a BFP reporter circuit in Jurkat T-cells via introduction of ortho-ligands that vary in their geometry and valency. **c)** Schematic of a modulation strategy for ortho-ligand responsive SNIPRs driving a BFP reporter circuit via introduction of ortho-ligands for a pseudo C2 structure by homodimerization. **d)** BFP reporter activation in Jurkat T-cells upon addition of C2 ligand and the OrthoTuner dimerization disrupter (0.05-500 nM) (N = 3; +/- SEM). **e)** Illustration demonstrating how private and promiscuous signaling channels were created by using a set of five heterodimer pairs (Left). These pairs were computational designed to have exclusive binding partners or accommodate promiscuous binding of other heterodimer components. (Right) Activation of five ortho-ligand responsive SNIPR driving a BFP reporter circuit in Jurkat T-cells via introduction of five ortho-ligands fused to a C6 scaffolding backbone. Ligands were added at 500nM concentration (N = 3; +/- SEM).

**Figure S2.**
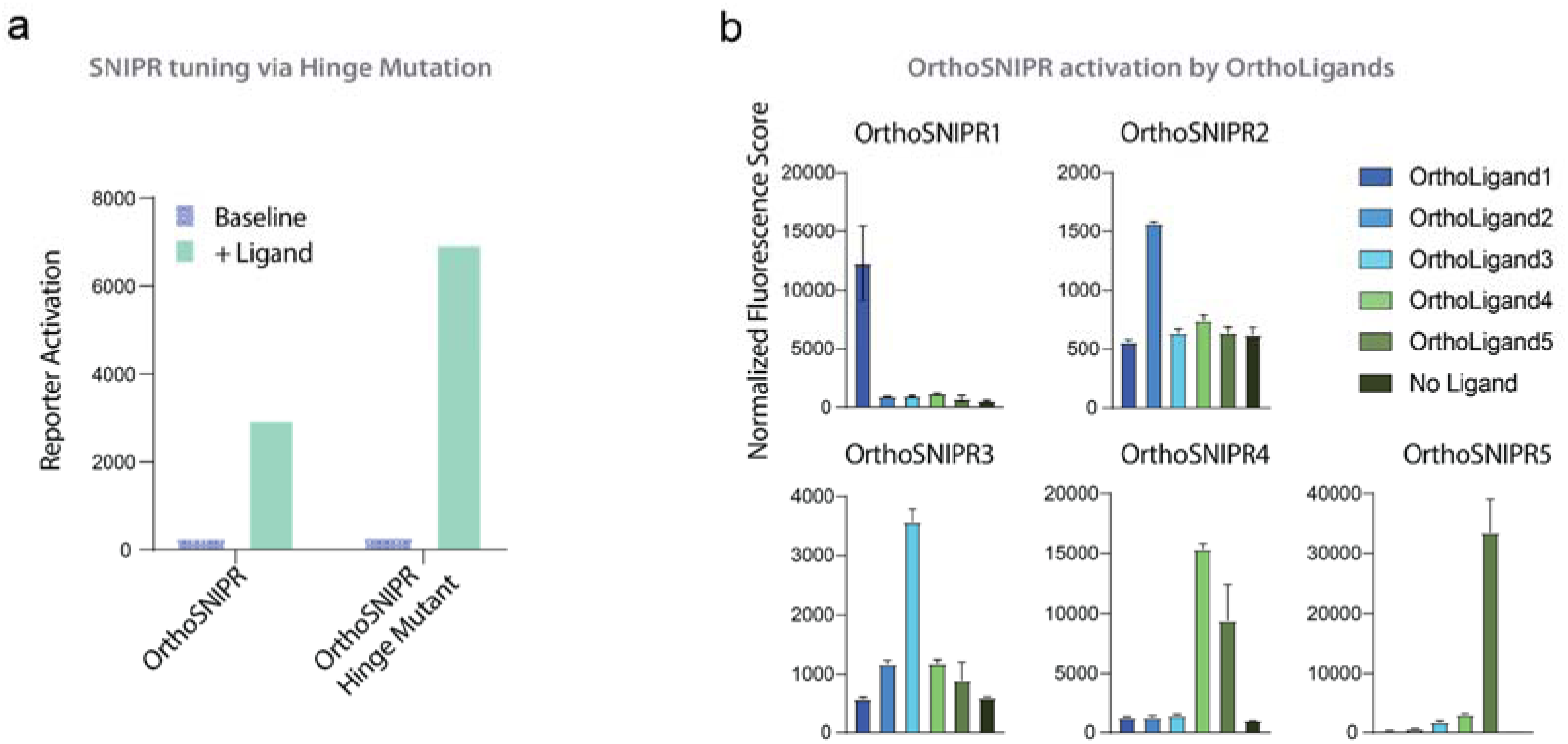
**a)** Activation of an orthogonal ligand responsive SNIPR or a modified SNIPR lacking the dimerization cysteine hinge residue via introduction of ortho-ligand C1-active at 200 nM (N = 1). **b)** Individual activation analysis of the 5×5 receptor ligand screen shown in Fig. 2e (N = 3; +/- SEM).

To enable the creation of bio-orthogonal communication networks, we generated a series of OrthoSNIPR receptor-ligand pairs using five different designed heterodimer pairs. As above, we utilized one subunit of each heterodimer as the extracellular domain of the SNIPR, and the second subunit was fused to a designed C6 scaffold conferring robust signaling. To maximize the versatility of the signaling networks that could be generated from these synthetic receptors, we chose three heterodimer pairs that previous characterization with purified proteins showed are completely orthogonal, and two heterodimers that cross interact; the former enable insulated communication pathways, the latter combinatorial modulation as observed with BMP signaling (**Figure 2e**, **Supplementary Figure 2b**). As predicted from the biochemical properties of the binding domains, three of the receptors are completely orthogonal and respond only to their ligands, while the other two recapitulate the more relaxed specificity of the parent designs. Such cross-reactive components enable the design of complex networks with fewer parts than would be possible with fully orthogonal receptors, allowing for encoding more information in therapeutic cells by payload-limited gene delivery vectors. Thus, the OrthoSNIPR system can be readily expanded beyond one receptor to form the basis of a complex communication network that relies on private and promiscuous signals to process and respond to their environments and execute user-defined functions in therapy, development, or homeostasis.

## SNIPRs Access a Distinct Activation Mechanism when Sensing Soluble Ligands

The previously undescribed capacity of SNIPRs to detect soluble ligands is a novel behavior for engineered Notch-like receptors. Understanding this mechanism of soluble factor-regulated SNIPR activation could have implications for further receptor engineering and optimization while potentially elucidating new possibilities for Notch signaling beyond what has been observed in nature. Notch and synNotch signaling is typically regarded as being restricted to surface-bound ligands^17^, and previous literature has explicitly demonstrated the inability of a soluble ligand to activate synNotch in contrast to a membrane-tethered analogue^7,18^. The homodimeric nature of TGF-β and VEGF suggested a model of mechanical cross-activation in trans, but SNIPR activation is insensitive to T-cell dilution (**Supplementary Figure 3a**). An alternative putative model proposed that ligands may apply force on the receptor by adsorbing to the culture plate, but using low-binding plastic vessels did not abrogate signaling (**Supplementary Figure 3b**).

Beyond the canonical mechanical model of force-mediated Notch signaling at the plasma membrane, evidence for the role of receptor endocytosis in Notch activation has emerged^19^, and the localization of γ-secretase activity to acidic compartments implies that pH may also play a role in some contexts^20^. To probe the behavior of SNIPRs as they enter the activation pathway, we perturbed each potential step using small molecule inhibitors of the cognate molecular process (**Figure 3a**). All Notch-based receptors from soluble SNIPRs to CD19-responsive synNotch are sensitive to γ-secretase inhibition via DAPT, implying a common final trigger for transcription factor release, while doxycycline-mediated induction from the control pTRE promoter was insensitive to this perturbation (**Figure 3b**). Surprisingly, we found that inhibition of ADAM protease, which is responsible for the proximal cleavage event upon Notch ligand binding and LNR conformational change, hindered the activation of both the CD19-responsive synNotch and SNIPR, but did not affect the soluble SNIPRs. In contrast, blocking endosomal acidification via chloroquine selectively inhibited the soluble SNIPRs, suggesting a model in which soluble factor binding triggers an endocytic cascade culminating in proteolysis within the endosomal membrane. The orthoSNIPRs bearing artificial soluble ligand:receptor pairs recapitulated this result. Inhibition of the TGF-ꞵ SNIPR by chloroquine was dose-dependent (**Supplementary Figure 3c**) while the effect of the DMSO vehicle was minimal at the assayed concentrations (**Supplementary Figure 3d**). In accordance with the receptor inhibition assay, confocal imaging confirmed that a fluorescently labeled orthoSNIPR ligand colocalizes with LysoTracker, an endocytic marker, upon ligand addition (**Figure 3d**). SNIPRs remain predominantly localized to the plasma membrane, suggesting that a small fraction of activated receptors is responsible for the observed transcriptional activation.

**Figure 3:**
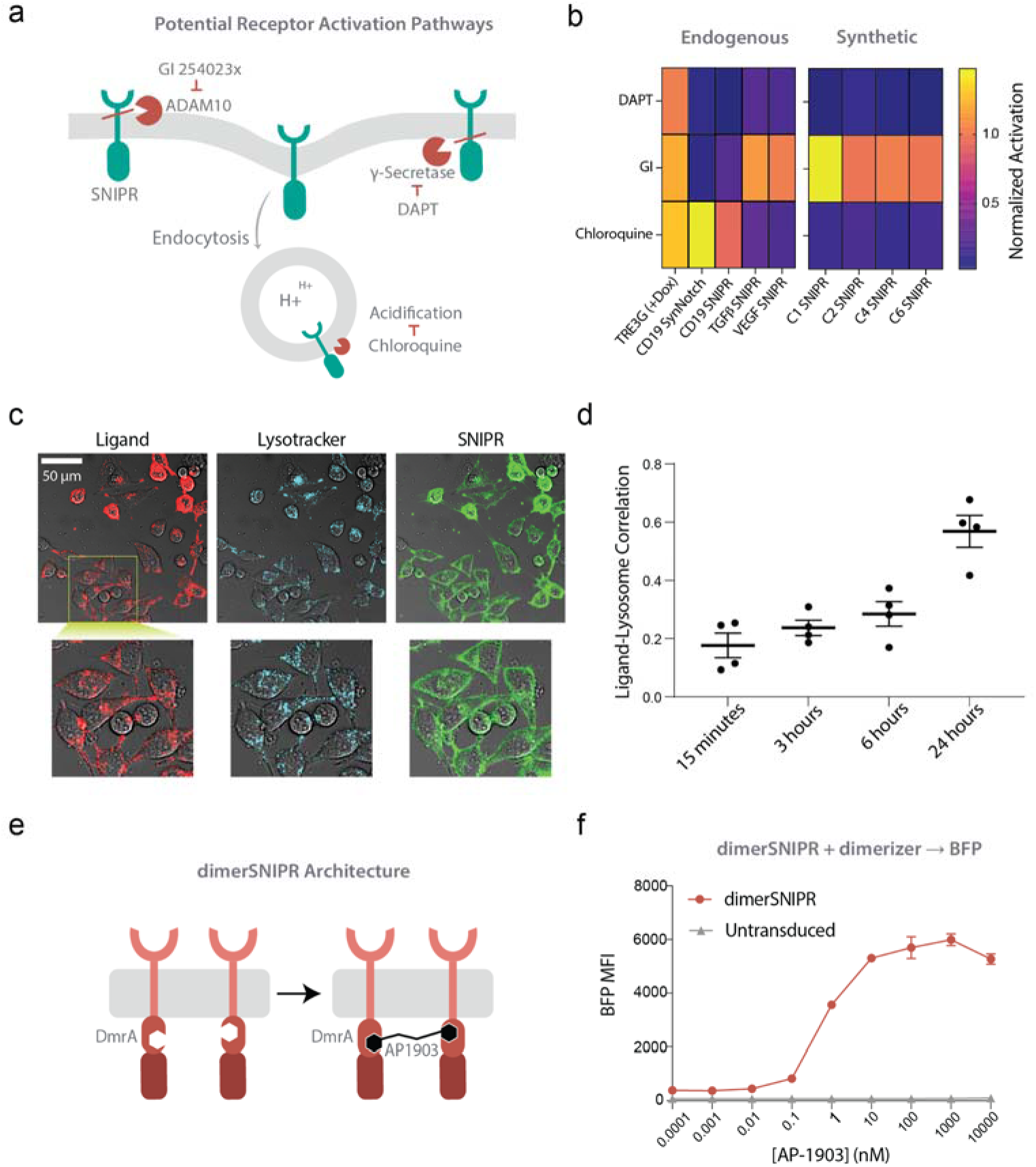
SNIPRs access a distinct activation mechanism. **a)** Schematic of potential activation pathways for soluble or conventional SNIPRs, along with chemical biology strategies to perturb these reactions. The conventional mechanism of proteolytically regulated receptors depends on ligand-mediated engagement of cell surface proteases (conventionally ADAM10, which is inhibited by the small molecule GI 254023x). The extracellularly truncated receptor becomes a substrate for y-Secretase, whose activity is blocked by DAPT. A proposed alternative pathway depends on endocytic activation of the receptor, where the terminal y-Secretase cleavage step may be mediated by endosomal acidification, which is inhibited by Chloroquine. **b)** Relative activation of primary human CD3^+^ T-cells by endogenous ligands, or Jurkat T-cells by synthetic ligands, in the presence of the chemical inhibitors described in **a**. T-cells expressing soluble or conventional SNIPRs, a conventional synNotch receptor, or an proteolysis-independent chemically-inducible promoter, were selected as a representative panel. Data represents a mean of N=3 measurements normalized to the activation of an inhibitor-free control sample. **c)** Confocal images of HeLa cells expressing a SNIPR fused to a GFP which were incubated with mCherry labeled C6-101A ligands for 24 hours and stained with Lysotracker marker for 30 minutes before imaging. **d)** Images were acquired at timepoints [15 minutes, 3 hours, 6 hours, and 24 hours] for the experiment in **(c)** and analyzed for colocalization between the mCherry labeled ligands and lysotracker signal. (N = 4 images per time point with at least 10 cells per image ; +/- SEM) **e)** Schematic of dimerSNIPR architecture. A DmrA homodimerizing protein domain was inserted into a TGF-β-responsive SNIPR between the juxtamembrane domain (JMD) and the transcription factor. The activation assay was performed in the absence of the extracellular TGF-β ligand. **f)** Activation of primary CD3^+^ T-cells via titration of the small molecule AP1903 (N = 2; +/- SD).

**Figure S3:**
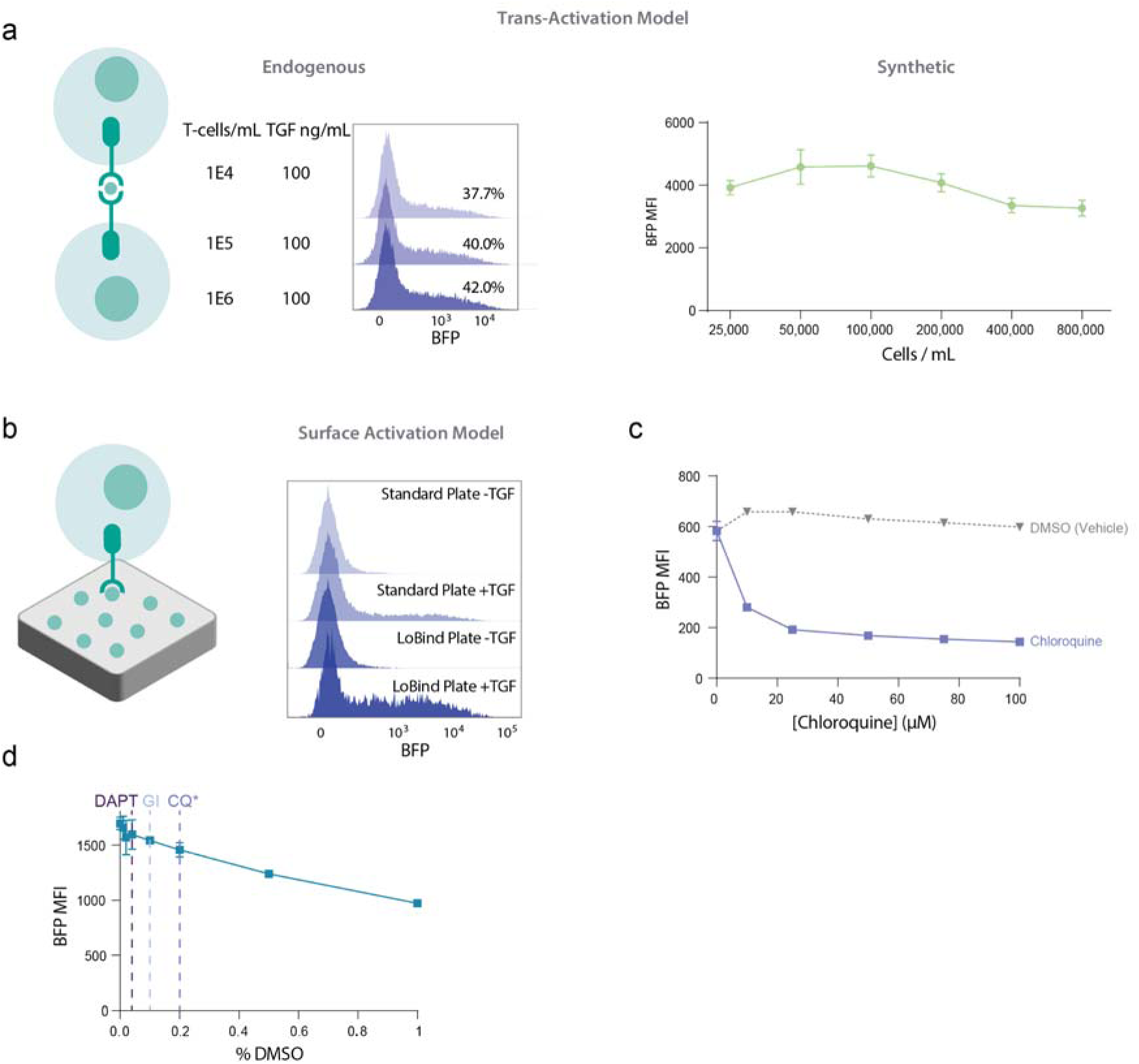
**a)** (Left) Schematic of a potential model of mechanosensitive soluble SNIPR activation wherein the mechanical force is applied via the simultaneous binding of a dimeric ligand by two SNIPR-expressing cells in trans. (Center) BFP reporter expression histograms depicting a T-cell titration assay wherein the TGF-β ligand concentration remains constant but the T-cells are diluted within the same volume. (Right) Activation of an ortho-ligand responsive SNIPR driving a BFP reporter circuit in Jurkat T-cells via introduction of ortho-ligand C1-active at 500nM while titrating down the number of receiver cells (N = 3; +/- SEM). **b)** (Left) Schematic of an alternative force-mediated soluble SNIPR activation mechanism wherein mechanical stress is applied via T-cell motion while the receptor binds to a putatively soluble ligand which has been immobilized on the cell culture vessel via adsorption. (Right) Representative histograms showing TGF SNIPR activation in the presence or absence of 50 ng/mL TGF-β1 in a conventional polystyrene vs. Low-Binding culture vessel. **c)** BFP reporter expression of TGF SNIPR T-cells upon addition of 100 ng/mL TGF-β and the indicated concentration of Chloroquine or vehicle control. **d)** BFP reporter expression of TGF SNIPR T-cells in the presence of 100 ng/mL TGF-B and the indicated concentration of DMSO. Vertical dashed lines indicated the final DMSO vehicle concentrations for DAPT, GI 254023x, and Chloroquine in **Figure 3b** (left). Note that the Chloroquine and vehicle concentrations used to assay the synthetic receptors were 4x lower than that in the endogenous receptor panel.

Despite the robust data suggesting SNIPR activation via the endosomal axis, the mechanism by which ligand binding could stimulate receptor internalization remained unclear. A potential mechanism was suggested by the observation that the SNIPRs that were successfully generated to respond to natural ligands all bind to dimers, similarly to orthoSNIPRs that bind to dimers or higher order oligomers. Receptor oligomerization may induce endocytosis via an avidity-like mechanism in which multiplexed low affinity binding motifs against the native endocytic machinery cooperatively drive internalization of the clustered complex^22^. We therefore hypothesized that ligand-mediated SNIPR dimerization may be responsible for receptor internalization. When the chemically-inducible DmrA (FKBP) homodimerization domain^23^ was inserted into the SNIPR scaffold (**Figure 3e**), the receptor demonstrated strong ligand-independent activation upon addition of the AP1903 homodimerizer (**Figure 3f**). Ligand-induced receptor multimerization may serve as the proximal trigger for the endocytic cascade culminating in endosomal transcription factor release and gene expression.

## Programming Cell:Cell Sensing and Responses with Soluble SNIPRs

To demonstrate that SNIPRs could detect physiological production of endogenous factors ligands rather than recombinantly expressed and purified ligands, we assayed SNIPR mediated BFP reporter expressing T-cells in co-culture with a panel of potential target human tumor cell lines (**Figure 4a**). We likewise extended our work on the orthoSNIPR system to demonstrate autonomous cell signaling with cell-derived rather than exogenous ligands. To this end, we engineered a “sender cell” that secretes the ligand. When the “receiver” T cells expressing the cognate receptor were cultured in the presence of the media conditioned by the sender cells, we observed signaling in response to the secreted ligand (**Figure 4b**). We chose to isolate the sender cells from the receiver cell to avoid cell membrane-based trans-activation and guarantee that any response is due to the soluble ligand. Sender cells secreting IGF-EndoTag1, an internalization-driving chimeric binding protein (see accompanying manuscript), strongly potentiated the SNIPR signaling. Our experiment demonstrated that with the orthoSNIPR communication system, cells can autonomously communicate with each other and share information in a channel that is completely private and bioorthogonal.

**Figure 4:**
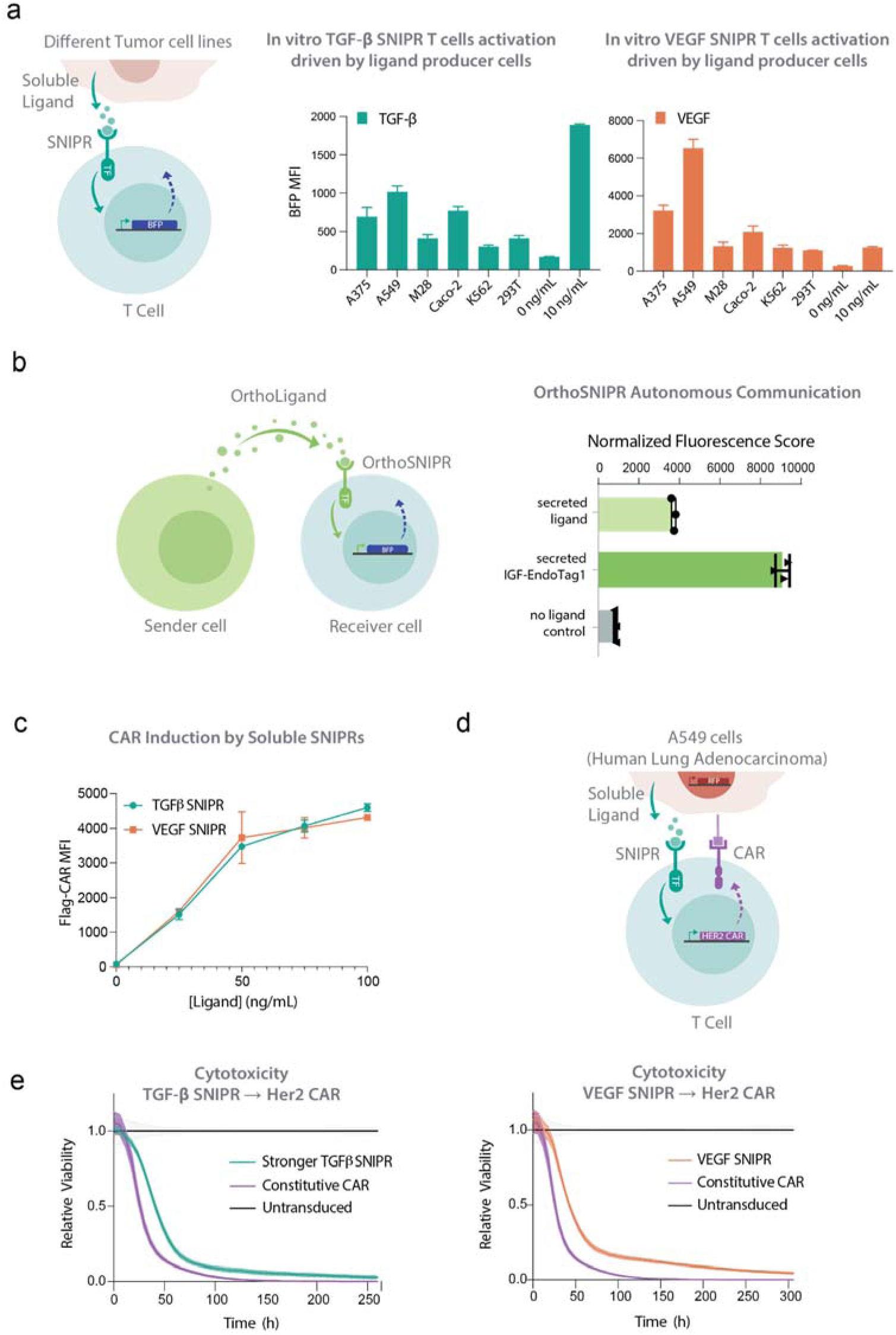
In vitro evaluation of engineered Cell-to-Cell communication via soluble ligands. **a)** (Left): Schematic of in vitro activation experiment. Co-cultured cancer cell lines secrete endogenous soluble factors which activate the cognate SNIPR, driving expression of the downstream BFP reporter. (Middle, Right): In vitro activation of primary human T-cells bearing TGF-β or VEGF-responsive SNIPRs in coculture with the respective cancer cell line (N = 3; +/- SD). **b)** Left: illustration of autonomous cell communication where a secretor cell releases an ortho-ligand into the extracellular environment and consequently activates receiver cells with a matching receptor. Right: Activation of an ortho-ligand responsive SNIPR driving a BFP reporter circuit in Jurkat T-cells via introduction of ortho-ligands which were secreted from another cell (N = 3; +/- SEM). **c)** Induction of a Flag-tagged HER2 CAR by TGF or VEGF SNIPR T-cells using titrated recombinant ligand (N=3; +/- SD). **d)** Schematic of in vitro cytotoxicity model. A549 lung adenocarcinoma cells transduced with a lentiviral vector encoding nuclear localized RFP to facilitate live cell imaging were co-cultured with primary human T-cells bearing a soluble SNIPR driving a HER2 CAR. **e)** (Incucyte) cytotoxicity assays of TGF-β (Left) or VEGF (Right) SNIPR circuit CD8^+^ T-cells, or constitutive HER2 CAR-T cells, targeting A549 cells. Cells were co-cultured in a 2:1 E:T ratio.

**Figure S4:**
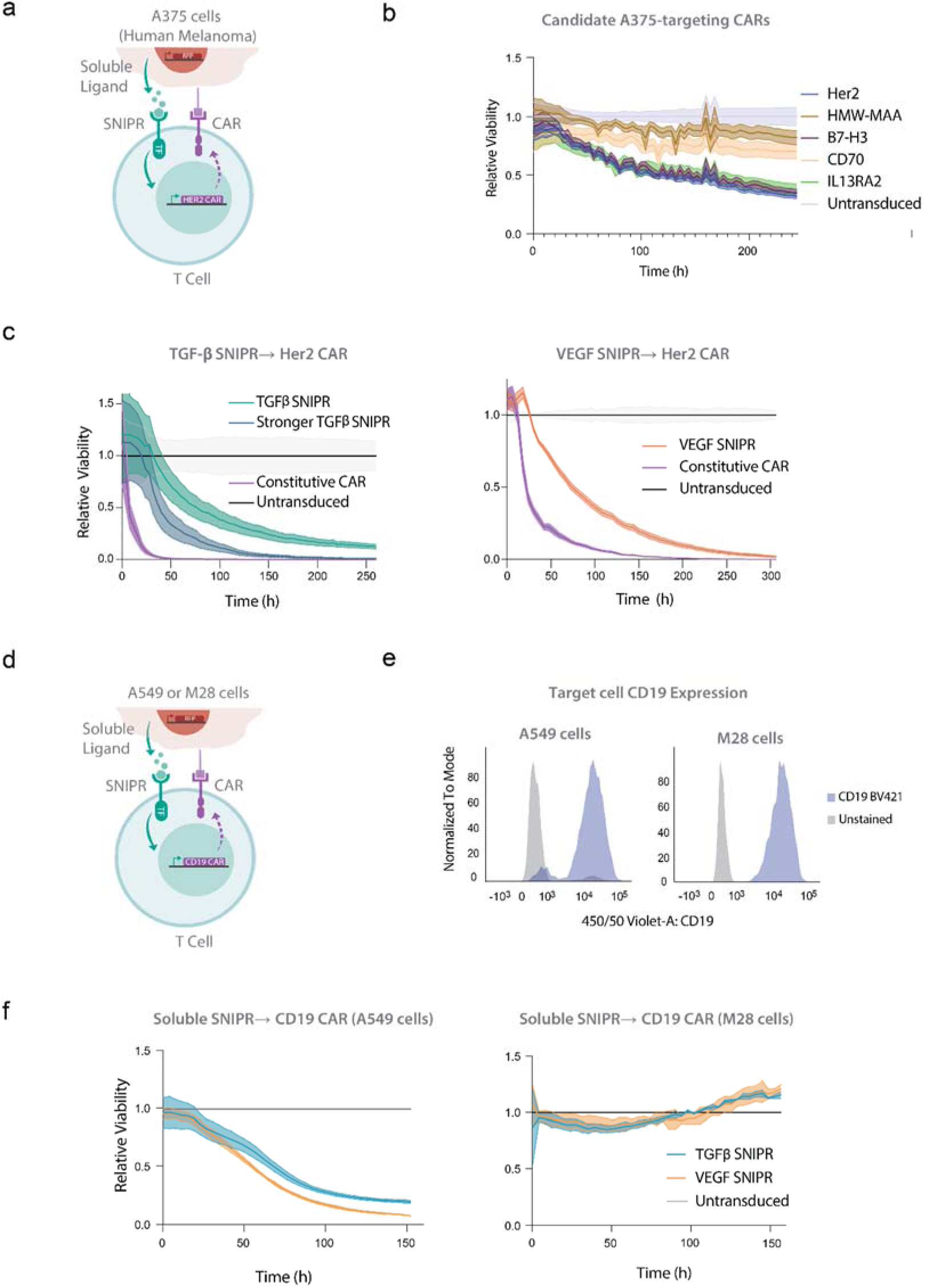
**a)** Schematic of cytotoxicity assay; RFP-labeled A375 melanoma cells induce production of a CAR via stimulation of a soluble SNIPR via endogenous TGF-β or VEGF production. **b)** Comparison of potential CAR targets endogenously expressed by A375 melanoma cells. CD8^+^ T-cells bearing a panel of TGF SNIPR → CAR circuits were incubated with RFP-labeled A375 cells and monitored via Incucyte imaging. **c)** Killing performance of Her2-directed CARs driven by TGF SNIPR variants (middle) or a VEGF SNIPR (right) vs. constitutive and untransduced controls. The experiment was performed via Incucyte live cell imaging. **d)** Schematic of in vitro killing specificity assay. High ligand-expressing (A549) and low-expressing (M28) target cells, as determined in **Figure 4b**, were transduced with a lentiviral vector encoding CD19 and cocultured with primary human T-cells bearing SNIPR → CD19 CAR circuits. **e)** Comparison of CD19 expression on the A549 (left) and M28 (right) target cells. **f)** Incucyte live cell imaging viability analysis of A549 (left) and M28 (right) cells ectopically expressing CD19 in the presence of TGF or VEGF-responsive SNIPR → CD19 CAR-T cells.

## Expanding the Landscape of Targetable Antigens for Engineered T cell Therapies with Soluble SNIPR **→** CAR Circuits

Soluble antigen sensors could improve the safety and efficacy of CAR-T therapy by selectively restricting CAR expression within the tumor, imparting asynchronous IF → THEN combinatorial logic and enhancing the specificity of potentially promiscuous CARs. A receptor intended for therapeutic use must strike a balance between low basal signaling to avoid off-target toxicity and robust activation to drive a sufficient level of CAR transgene expression for effective cancer cell killing. This is particularly challenging given the lower sensitivity of CARs compared to natural TCR counterparts^24^. We sought to construct circuits in which soluble SNIPR-primed activation drives expression of a CAR capable of addressing solid tumor antigens (**Supplementary Figure 4a**). To determine the optimal CAR payload for targeting A375 melanoma cells, a validated TGF-β producing solid tumor line, we first constructed TGF-β SNIPR → CAR circuits bearing a panel of CARs against potential endogenous A375 antigens and assayed their in vitro killing efficacy using a live cell imaging assay (**Supplementary Figure 4b**). Of the variants tested, a Her2-specific CAR was the most efficacious and was therefore selected for subsequent experiments.

Effective tumor clearance by CAR-T cells requires robust CAR expression. We verified that, similarly to BFP, the CAR payload is expressed by SNIPR → CAR T-cells in a dose-dependent manner (**Figure 4c**). We then showed that both TGF-β and VEGF-responsive SNIPR circuits driving the Her2 CAR exhibit robust killing of both A549 lung adenocarcinoma cells (**Figure 4d**, **e**) and A375 cells (**Supplementary Figure 4c**). Consistent with previous work, SNIPR → CAR circuits displayed slower killing kinetics than constitutively expressed CAR, owing to the temporal delay in CAR expression subsequent to primary ligand encounter^9^. To verify that target cell killing is not due to non-specific leaky CAR expression, we assayed the performance of SNIPR → CD19 CAR circuits against M28^CD19+^ cells (**Supplementary Figure 4d, e**), which only weakly stimulated the TGF-β and VEGF SNIPRs, and found that these cells resist killing (**Supplementary Figure 4f**).

We next evaluated the in vivo performance of SNIPR → CAR circuits. Mice bearing subcutaneously embedded A375 melanoma tumors underwent adoptive T-cell transfer 7 days post tumor seeding. TGF-β SNIPR → Her2 CAR T-cells significantly slowed tumor growth relative to untransduced control T-cells, while SNIPRs bearing the activating hinge mutation eliminated the tumors with efficacy similar to that of the constitutive CAR (**Figure 5a**). To assess the ability of soluble SNIPRs to improve the therapeutic window of CARs prone to off-tumor toxicity, we substituted the human-specific 4D5 Her2 CAR for a recently described mouse/human cross-reactive DARPin-based variant^25^. Constitutive DARPin CAR-T cells administered to NSG mice demonstrate toxicity and weight loss due to on-target/off-tumor toxicity against low levels of Her2 expression in lung tissue^26^, consistent with the fatal pulmonary toxicity reported in an early clinical trial of human Her2-targeting CARs^27^. SNIPR → CAR circuits are expected to enhance the safety of CARs with such promiscuous activity by confining their expression to the tumor (**Figure 5b**). The ability of TGF-β and VEGF SNIPRs to activate in response to both human and mouse target ligands (**Supplementary Figure 5a**) further enhances the clinical relevance of this model, although TGF-β and VEGF expression in immunocompromised NSG mice may not recapitulate that of syngeneic models.

**Figure 5:**
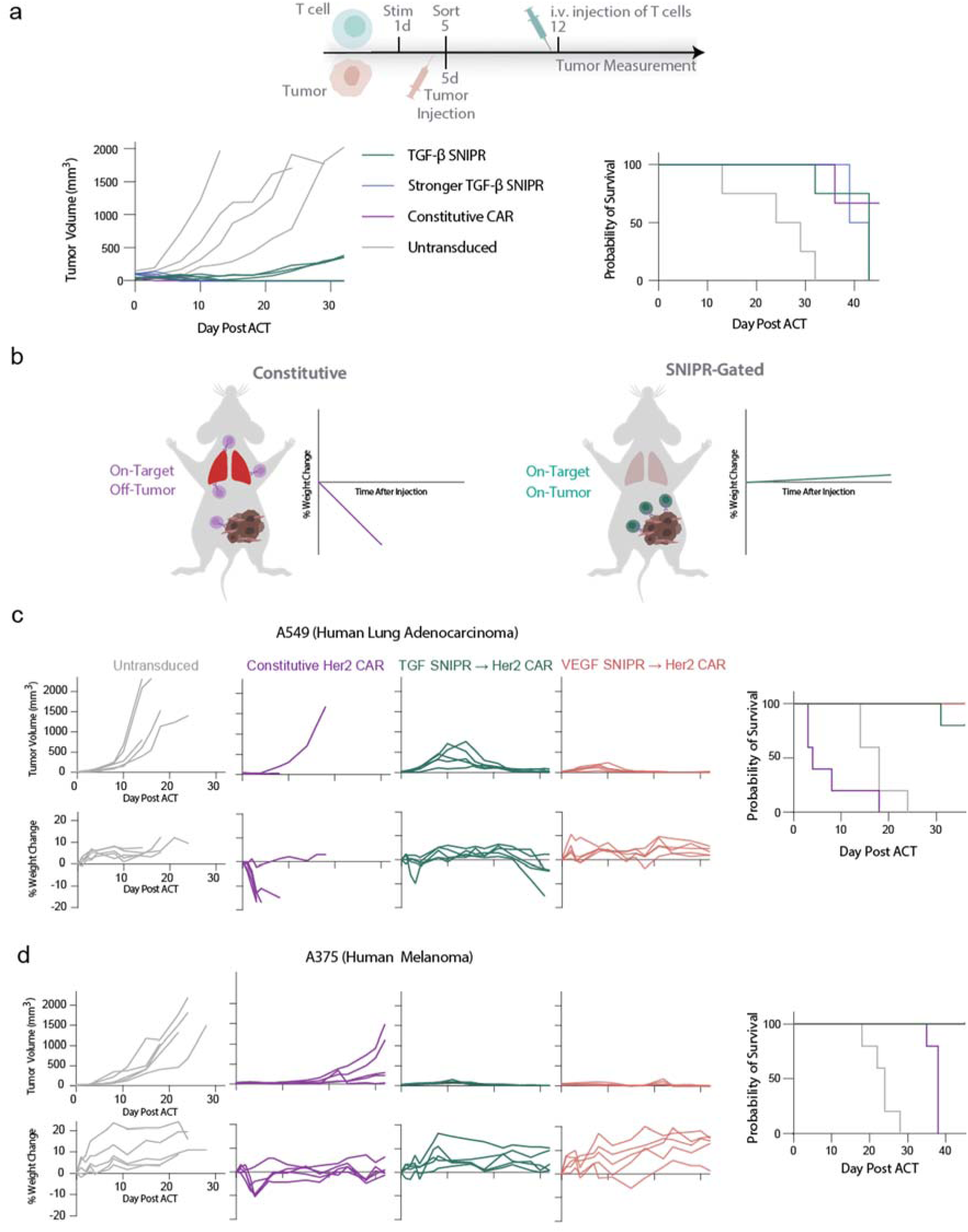
In vivo anti-tumor efficacy of Soluble SNIPR → CAR T-cells. **a)** NSG mice were implanted subcutaneously with A549 lung carcinoma or A375 melanoma cells. After 7 days, mice were adoptively transferred with primary human CD3^+^ T-cells bearing soluble SNIPRs driving a mouse/human cross-reactive Her2 CAR, the same CAR in a constitutive expression vector, or an untransduced control. Tumor volume (left) and Kaplan-Meyer survival plots (right) are shown. **b)** Schematic of in vivo CAR efficacy and toxicity study. The cross-reactive CAR has previously been shown to induce toxicity, most prominently displayed as a weight loss phenotype, due to On-Target/Off-tumor activity primarily targeting lung tissue. **c)** Tumor volume (top) and weight change (bottom) of A549 xenografted mice treated with untransduced, constitutive, or soluble SNIPR-gated expression of the cross-reactive Her2 CAR. Cumulative mouse survival is shown on the right. **d)** Activity of SNIPR circuits driving the ^tr^DARPin Her2 CAR in A375 melanoma xenografted mice is presented analogously to the data shown in **c**.

**Figure S5:**
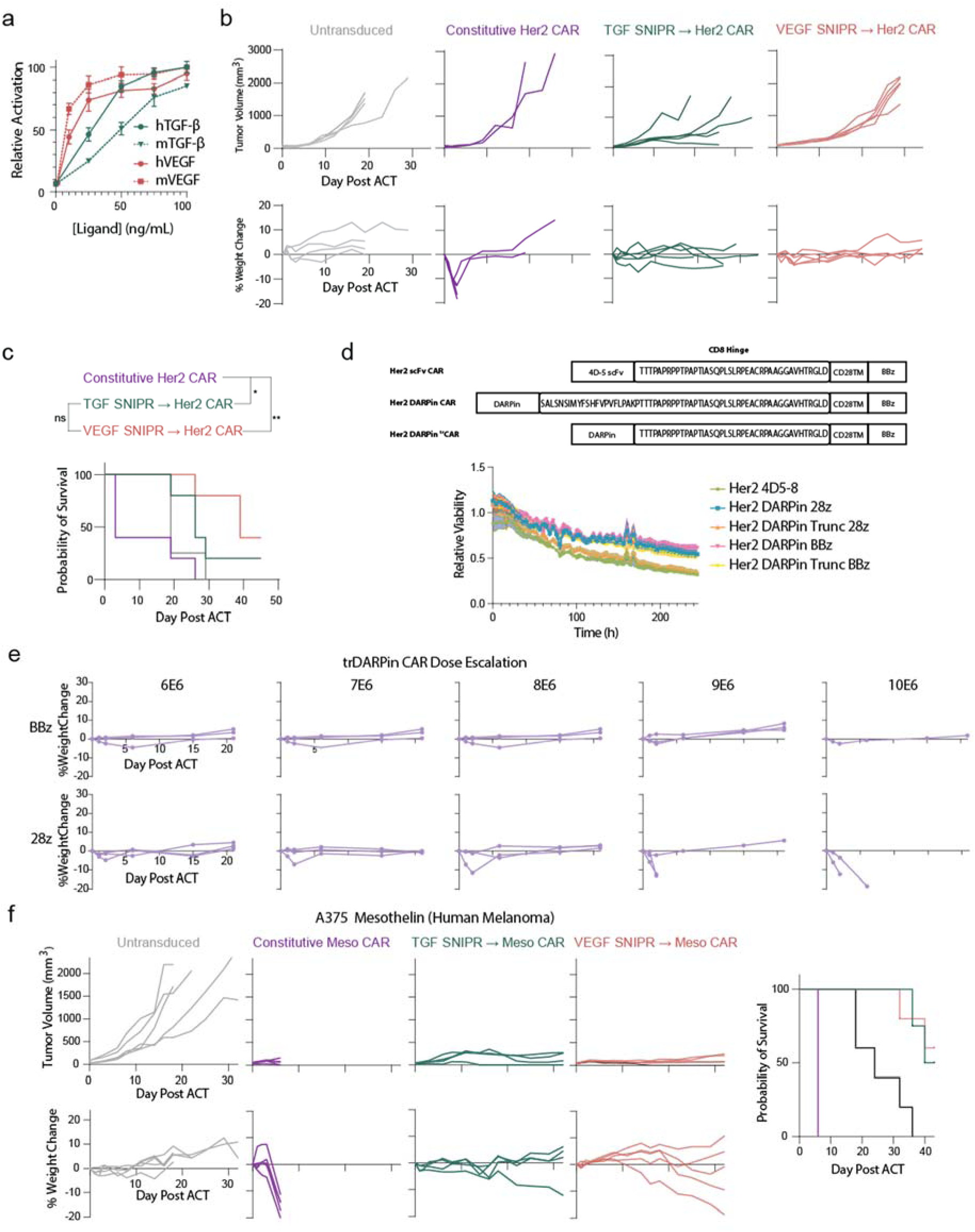
**a)** Comparison of TGF-B (left) and VEGF (right) SNIPR reactivity to recombinant human and mouse cognate ligands. Activation was assessed via induction of a BFP reporter gene (N=3; +/- SD). **b)** Tumor growth (top) and weight change (bottom) of NSG mice bearing A375 xenografts treated with untransduced, constitutive, or SNIPR → Her2 DARPin CAR-T cells. **c)** Kaplan Meyer survival curve for mice shown in **b**. **d**) Top: schematic of CD8 hinge sequence variation in the original 4D-5 CAR, original DARPin CAR, and matched truncated DARPin CAR. Bottom: comparison of cytotoxic potential of HER2 DARPin CARs in original or truncated CD8 Hinge architectures and bearing CD28 vs. 4-1BB costimulatory domains. CD8^+^ circuit-expressing T-cells were incubated with A375^RFP^ target cells and target cell survival was assayed via Incucyte imaging. **e)** Weight loss phenotype of tumor-free NSG mice injected with constitutive ^tr^DARPin Her2 CAR-T cells bearing 4-1BB (top) or CD28 (bottom) costimulatory domains. **f)** In vivo evaluation of a constitutive or SNIPR-gated mouse/human cross-reactive Mesothelin CAR in an NSG xenograft model. Tumor volume (top), weight change (bottom), and overall mouse survival (right) are shown.

We first evaluated the performance of our SNIPR → DARPin CAR circuits in an A549 adenocarcinoma xenograft model. Consistently with the previously published data, mice injected with constitutive DARPin CAR-T cells experienced rapid weight loss to the humane euthanasia threshold (**Figure 5c**). In contrast, mice receiving the SNIPR → CAR circuits maintained a healthy weight and were able to robustly control tumor growth, emphasizing the potential of this approach for generating potent cell-based therapies with improved safety profiles. We then sought to validate our circuits in an A375 melanoma xenograft model. Despite retarding tumor growth, the SNIPR circuit T-cells failed to clear the tumors, suggesting that A375 xenografts are more resistant to T-cell cytotoxicity (**Supplementary Figure 5b, c**). In vitro cytotoxicity analysis revealed that the DARPin CAR, constructed as reported with an extended CD8 hinge domain, is less potent than the original 4D5-based version, but we found that a modified construct bearing a truncated hinge matching that in the 4D5 receptor (DARPin ^tr^CAR) partially restored efficacy (**Supplementary Figure 5d**). Unexpectedly, we also found that this truncation reduced in vivo toxicity of the constitutive CAR-T cells, necessitating the reinstallation of the CD28 costimulatory domain and injection of a higher dose to recapitulate weight loss (**Supplementary Figure 5e**).

We then evaluated the performance of the SNIPR → ^tr^CARs in the A375 xenograft model, increasing the tumor dosage proportionally to the T-cell amount. The TGF-β and VEGF SNIPR → ^tr^CAR circuits were well-tolerated and highly efficacious, while constitutive DARPin ^tr^CAR-T cells rapidly induced weight loss within several days of infusion, albeit not to the euthanasia threshold, and then failed to control tumor outgrowth (**Figure 5d**). To test an orthogonal toxicity model, we also evaluated our constructs using a cross-reactive anti-Mesothelin nanobody-based CAR payload^28^ using A375 tumor cells ectopically expressing Mesothelin (**Supplementary Figure 5f**). Similarly to the cross-reactive Her2 CARs, the SNIPRs significantly increased the therapeutic window of this potent and highly toxic CAR.

## Discussion

The SNIPR architecture satisfies the demands of high performance soluble factor sensing. SNIPRs bearing ligand binding domains targeting cancer-associated factors show robust activation upon titration of recombinant ligands, and are capable of driving potent therapeutic responses at the site of disease mitigating toxicity potential of cell therapies. The ability of SNIPRs to recognize engineered orthogonal ligands expands their potential capabilities beyond natural cues and unidirectional signaling. Immune cells bearing orthoSNIPRs could coordinate to establish signaling feedback loops to more precisely pinpoint tumor coordinates within the body, and different immune cell types bearing their own receptors and payloads could interact to activate specific programs at the optimal times during treatment stages. Customization of orthoSNIPR signaling using privileged or promiscuous signaling channels enables fine tuning of engineered cell behavior, while the capability to sense and secrete endogenous signaling factors provides an interface with the local host immune environment.

Overall, soluble SNIPRs provide a versatile tool for therapeutic bioengineering and other disciplines in biology. The customizability of these receptors could enable them to sense morphogen gradients established by proteins such as NGF and the BMP family during embryonic development, or report on the local immune state in the context of cancer, autoimmunity, and infectious disease. These receptors are also suitable for use in developing complex high-order biocomputational circuits owing to their programmability via synthetic ligands and compatibility with modular payloads. The compact genetic footprint may permit the development of multi-receptor circuits that integrate various soluble or cell surface inputs for precise localization of biological activity^29^. SNIPRs are an expanding platform for precise control of cells for therapeutic purposes and basic biology applications.

## Methods

### Plasmid Assembly

Vectors were derived from previously reported methods, with receptor vectors originating in pHR_PGK (AddGene 79120) and reporter constructs cloned into pHR_Gal4UAS_PGK_mCherry (AddGene 79124). All constructs reported in this work were generated via NEBuilder HiFi DNA Assembly (NEB E2621). A mapping of which constructs correspond to which figures is shown in **Supplementary Table 1**. Sequences are listed in **Supplementary Table 2**. The origins of new components described in this manuscript are listed in **Supplementary Table 3**.

### Cell culture and lentiviral production

Lenti-X 293T cells (Clontech 632180) were cultured in DMEM supplemented with 10% FBS, 1x Sodium Pyruvate (Millipore Sigma), and 50 U/mL penicillin-streptomycin. A375 cells expressing nuclear-localized RFP were a generous gift of the Marson lab at UCSF. A549 cells (CCLZR013) were obtained from the UCSF Cell and Genome Engineering Core. A375 and A549 cell lines were grown in 293T media. Caco-2 cells were obtained from the ATCC (HTB-37) and grown in DMEM (Gibco 11965092) supplemented with 20% FBS and 50 U/mL penicillin-streptomycin. M28 cells were obtained from the Gerwin laboratory at the National Cancer Institute and grown in RPMI 1640 (Gibco) with 10% FBS, 50 U/mL penicillin-streptomycin, and 1x GlutaMAX (Gibco). K562 cells (ATCC #CCL-243) were cultured in IMDM (Gibco) with 10% FBS and 50 U/mL penicillin-streptomycin. For viral production, 1E6 Lenti-X 293T cells were seeded in a 6 well vessel at 1E6/well in 2.5 mL of 293T media. One day after seeding, cells were transfected with a packaging mix consisting of 1.5 μg of the transgene expression vector, 1.34 μg of pCMVdR8.91, and 0.17 μg of pMD2.G, using TransIT-Lenti Transfection Reagent (Mirus) (3 μl of the reagent per 1 μg of total DNA). Two days after transfection, viral supernatants were harvested via centrifugation and used immediately for transduction.

For Jurkat experiments, HEK293T cells () were initially cultured in DMEM (Gibco) supplemented with 10% FCII (Gibco), and 50 U/mL penicillin-streptomycin (Gibco), then with “induction” DMEM media supplemented with 10% FCII, 10 mM sodium butyrate, and 50 U/mL penicillin-streptomycin, and later with “viral” DMEM media supplemented with 10% FCII, 0.1 mM HEPES, 1x GlutaMax, 1x MEM non-essential amino acids () and 50 U/mL penicillin-streptomycin. Jurkat T Cells were cultured in RPMI 1640 supplemented with 1x GlutaMax (Gibco), 10% FCII, and 50 U/mL penicillin-streptomycin. For lentivirus production, 293T cells were seeded at 400k cells/mL in either a 12-well vessel or T75 plate and grown to 90% confluency. Then, 22 ug of expression vector was mixed with 22 ug of pCMV and 4.5 ug of pVSV-G (for 12-well transfections these ug quantities of DNA were divided by 4). This DNA mixture was then packaged using PEImax () and 1x OptiMem (Gibco) and added to the media of the 293T cells. Then, 12-17 hours later the media was removed from these cells and replaced with “induction” DMEM media containing 10 mM sodium pyruvate. After incubating for 6-8 hours, the “induction” media was replaced with “viral” media. Two days later, viral supernatants were harvested using LentiX Concentrator (), followed by centrifugation and used immediately for transduction.

### Primary human T cell culture

T-cells were isolated from anonymized donor blood post-apheresis (Vitalant) either in bulk CD3^+^ format via positive selection (STEM CELL 17851) or as separated CD4^+^ or CD8^+^ populations (STEM CELL 15062 and 15063). Use of donor material was approved by the UCSF Institutional Review Board. Upon isolation, T-cells were frozen in liquid nitrogen in RPMI-1640 (Thermo Fisher 11875093) supplemented with 20% human AB serum (Valley Biomedical Inc., HP1022) and 10% DMSO. For experiments, T-cells were thawed at 37°C, washed, and cultured in human T-cell media consisting of X-VIVO15 (Lonza #04-418Q) with 5% Human AB Serum, 10 mM N-acetyl L-Cysteine (Sigma-Aldrich #A9165) neutralized with 1M NaOH (Sigma-Aldrich S2770) and supplemented with 30 units/mL of IL-2 (NCI BRB Preclinical Repository). One day post-thaw, T-cells were stimulated with washed Dynabeads (Thermo Fisher 11132D) using a 1:3 cell:bead ratio. On the following day, untitered lentiviral supernatant was added at a 1:1 total volume ratio. After 24 hours of infection, cells were gently pelleted (400xg for 5 minutes) and depleted media was exchanged for fresh complete media. After three days of subsequent expansion, Dynabeads were removed via magnetic separation and cells were sorted (Beckton Dickinson FACSARIA II) for populations expressing both epitope-tagged receptors and constitutive fluorophore-expressing reporter circuits. Experiments typically commenced 4 to 7 days post-sort. All experiments were performed in complete human T-cell media aside from Incucyte assays, which were performed in RPMI / 10% FBS.

### Flow Cytometry and Sorting

For sorting, T-cells were pelleted and resuspended in 100 μL of PBS with 2% FBS and antibody at a 1:100 ratio. After 20-30 minutes of staining at room temperature, cells were pelleted and resuspended in PBS/2%FBS and kept on ice until sorting. For flow cytometry measuring a fluorescent reporter gene, cells were directly analyzed and gated on scatter (FSC-A vs. SSC-A), single cells (FSC-H vs. FSC-A), and the constitutive co-fluorophore expressed by the reporter construct. For cytometry experiments involving immunohistochemistry, cells were washed once in PBS prior to staining for 20-30 minutes at room temperature, and then washed 3x prior to analysis. Antibodies used are listed in **Supplementary Table 4**.

### In Vitro SNIPR activation

Recombinant soluble factors were gently resuspended in their respective manufacturer-recommended reconstitution buffers and frozen at -80°C as single use aliquots. After thawing at room temperature, proteins were diluted in T-cell media to the appropriate concentration and either added directly to the T-cells in 96 well plates at a 1:100 dilution, or pre-diluted in media to 2x final concentration and added in a 1:1 ratio. Cells were mixed briefly and incubated at 37°C for 24-72 hours as indicated, at which point reporter expression was assayed via flow cytometry (BD FacSymphony X50 SORP or LSR II SORP). For inhibitor experiments, DAPT, GI 254023X and Chloroquine were used at a working concentration of 10 μM, 10 μM and 25-100 μM, respectively. Recombinant soluble factors employed in experiments are listed in **Supplementary Table 5**, and inhibitors are shown in **Supplementary Table 6**. Where unspecified, TGF-β refers to human TGF-β1 and VEGF refers to human VEGFɑ. For the TRE3G promoter, doxycycline hyclate (ab141091) was added as the ligand analogue reagent.

For Jurkat experiments, Jurkat T Cells were harvested and resuspended at 400k cells/mL in fresh RPMI media. De-novo designed OrthoLigands were diluted to the appropriate concentrations in 1x PBS and were added to freshly washed cells at 1/10 dilution (20 uL ino 200 uL) in 96 well plates. Assay plates were incubated at 37 C for 17-24 hours and then analyzed using flow cytometry (Invitrogen Attune). The OrthoLigands and inhibitors used in these experiments are also listed in **Supplementary Table 5** and **Supplementary Table 6** respectively.

### Protein expression and purification

OrthoSNIPR ligands were expressed in E. coli BL21 (NEB). Briefly, the DNA fragments encoding the design sequences were assembled into pET-29 vectors via Gibson assembly and further transformed into BL21 strain with heat-shock. Protein expression was induced by the autoinduction system and proteins were purified with immobilized metal affinity chromatography (IMAC). The elutions were further purified by FPLC SEC using Superdex 75 10/300 GL or Superdex 10/300 200 columns (GE Healthcare). Protein concentrations were determined by NanoDrop (Thermo Scientific) and normalized by extinction coefficients. Proteins were diluted to the appropriate concentrations in 1x PBS and applied to cell culture media using 1/100 dilutions.

### Confocal microscopy

HeLa cells were engineered to express the SNIPR receptor with a cleavable GFP domain in place of a transcription factor. These cells were seeded in 18 well glass bottom µ-Slides (Ibidi, Cat. No. 81817) at a density of 15k/well. mCherry fused SNIPR ligands were incubated with the cultured cells for 15 min, 3 h, 6 h or 24 h,Lysotracker (ThermoFisherScientific, Cat. No. L7526) was added for 30 min before imaging. Cells were washed 3× in PBS and immediately proceeded to imaging. Confocal laser scanning microscopy was performed on a Nikon A1R HD25 system equipped with a LU-N4 laser unit (Lasers used: 488 nm, 561 nm, 640 nm). Data was acquired using an 20×, NA 0.75, WD 1.00 mm air objective (Plan Apochromat Lambda) in combination with 1 multialkaline (EM 650 LP) and 2 GaAsP detectors (DM 560 LP EM 524/42 (503-545) and DM 652 EM 600/45 (578-623)). Acquisition was controlled via NIS Elements software and data was analyzed via Fiji and custom-written Python Scripts.

### In vitro Cell-Cell communication assays

For in vitro detection of cell-produced TGF-β and VEGFɑ, target cells were seeded in a flat bottom 96 well culture vessel at 20k/well in each cell type’s native media. After 24 hours of adherence and growth, the growth media was aspirated and replaced with 40k/well T-cells in human T-cell media. BFP reporter activation was determined via flow cytometry after 48 hours of coculture.

To generate OrthoLigand sender cells, HeLa cells were transiently transfected via electroporation with a plasmid encoding an Ortholigand that was preceded by a modified serum albumin secretion tag. The cells were then grown for 48 hours to allow for protein expression before collecting the media. To eliminate any potential cell contamination, the media was centrifuged at 500xg. Receiver Cell Assay To assess the functionality of the Ortholigand, the conditioned media from the sender cells was added to Jurkat cells, which served as the receiver cells. The cells were incubated for 24 hours before being analyzed for BFP expression. BFP signal was indicative of successful Ortholigand-mediated signaling between the sender and receiver cells.

### Incucyte Imaging

For in vitro cell live cell imaging killing assays, A375, A549, or M28 target cells bearing nuclear-localized mKate2 were seeded in flat bottom 96 well plates in their native media. After 24 hours, media was aspirated and immediately replaced with T-cells in RPMI1640 (Gibco) + 10% FBS / + 50 U/mL penicillin/streptomycin + 30 U/mL IL-2 so as to maintain an expected 2:1 effector:target ratio at the start of the experiment. Plates were imaged using the Incucyte S3 Live-Cell Analysis System (Essen Bioscience) with 3 or 4 images collected per well and an imaging frequency of 4 - 6 hours for at least 7 days of coculture. Cells were counted via automated segmentation of the fluorescent target cell nuclei using the Incucyte software.

### In Vivo Xenograft models

All animal work was conducted under approval from the UCSF Institutional Animal Care and Use Committee (protocol # AN177022-03C). All experiments employed NOD.Cg-*Prkdc^scid^Il2rg^tm1Wjl^*/SzJ (NSG) (RRID:IMSR_JAX:005557) mice of age 8 - 12 weeks at the onset of experimentation. For all tumor models, dissociated cancer cells in 0.1 mL serum-free DMEM were implanted subcutaneously in the flank. For the A375 model targeting HER2, 1.5E6 tumor cells were injected. In all other experiments, 1E6 tumor cells were administered. T-cells were administered retro orbitally in 0.1 mL PBS 7 days post-tumor injection (5 days post-sort). For the A375 model targeting HER2, 9E6 T-cells were delivered; all other experiments used 6E6 T-cells per mouse. Tumor size was monitored via caliper, and for experiments employing mouse/human cross-reactive CARs, mice were weighed to a precision of 0.1 g daily for one week post T-cell injection and then at all subsequent tumor measurement time points. Mice were euthanized upon tumor measurement along any axis of > 20 mm or upon reaching a tumor volume of > 2000 mm^3^, where volume = ½ * largest axis * (smallest axis)^2^, or upon weight loss of 15% below initial weight at tumor injection as specified in our IACUC-approved protocol.

### Statistical Analysis

Data are presented as means +/- SD of technical replicates unless indicated otherwise in figure captions. For comparison of survival data, the two-tail P value for log-rank is reported.

### Software

Chart plotting and statistical analysis were performed in GraphPad Prism 9. Flow cytometry and sorting were performed using BD FACSDiva, and post-hoc gating and analysis were conducted using FlowJo 10.8.0. Where not visible in charts, error bars are smaller than symbols. Imaging colocalization analysis was performed with a custom-written python script based on the Pearson correlation coefficient.

## Author Contributions

D.I.P. and M.H.A. conceived and designed the initial project, interpreted data, and wrote the manuscript. M.J.D designed and performed experiments, interpreted data, and wrote the manuscript. A.C. performed experiments and interpreted data. I.Z. designed and performed experiments related to the IFN-γ SNIPR. P.T.R. designed, performed, and interpreted initial experiments. T.S performed experiments and interpreted data relating to cell imaging. B.H assisted with EndoTag ligand design and generation. D.L performed experiments relating to orthogonal ligand secretion. D.B. and K.T.R. supervised experiments and reviewed and edited the manuscript.

## Data Availability

All data associated with this study are present in the manuscript or its Supplementary Information files.

## Competing Interests

D.I.P., I.Z., and K.T.R. have patents for SNIPR receptors or variants. D.I.P. is an employee of Dispatch BioTherapeutics. K.T.R. reports personal fees from Arsenal Bio, Dispatch BioTherapeutics, Judo Therapeutics, and Alaunos Therapeutics.

## Description of Supplementary Materials

**Supplementary Table 1**: Plasmid names and T-cell donor information for each experiment described.

**Supplementary Table 2:** Description, part, and amino acid & DNA sequences for all plasmids described.

**Supplementary Table 3:** Description and sourcing of genetic components used in manuscript.

**Supplementary Table 4:** List of antibodies used in manuscript.

**Supplementary Table 5:** List of recombinant soluble factors for all experiments described.

**Supplementary Table 6**: List of pharmacological compounds used in receptor inhibition experiments.

## References

1. Schwank, G. & Basler, K. Regulation of Organ Growth by Morphogen Gradients. Cold Spring Harb. Perspect. Biol. 2, a001669 (2010).

2. Berraondo, P. et al. Cytokines in clinical cancer immunotherapy. Br. J. Cancer 120, 6–15 (2019).

3. Schwarz, K. A., Daringer, N. M., Dolberg, T. B. & Leonard, J. N. Rewiring human cellular input–output using modular extracellular sensors. Nat. Chem. Biol. 13, 202–209 (2017).

4. Kroeze, W. K. et al. PRESTO-Tango as an open-source resource for interrogation of the druggable human GPCRome. Nat. Struct. Mol. Biol. 22, 362–369 (2015).

5. Kipniss, N. H. et al. Engineering cell sensing and responses using a GPCR-coupled CRISPR-Cas system. Nat. Commun. 8, 1–10 (2017).

6. Mahameed, M., Wang, P., Xue, S. & Fussenegger, M. Engineering receptors in the secretory pathway for orthogonal signalling control. Nat. Commun. 13, 7350 (2022).

7. Morsut, L. et al. Engineering Customized Cell Sensing and Response Behaviors Using Synthetic Notch Receptors. Cell 164, 780–791 (2016).

8. Roybal, K. T. et al. Engineering T Cells with Customized Therapeutic Response Programs Using Synthetic Notch Receptors. Cell 167, 419–432.e16 (2016).

9. Zhu, I. et al. Modular design of synthetic receptors for programmed gene regulation in cell therapies. Cell 185, 1431–1443.e16 (2022).

10. Sloas, D. C., Tran, J. C., Marzilli, A. M. & Ngo, J. T. Tension-tuned receptors for synthetic mechanotransduction and intercellular force detection. Nat. Biotechnol. 1–9 (2023) doi:10.1038/s41587-022-01638-y.

11. Chang, Z. L. et al. Rewiring T-cell responses to soluble factors with chimeric antigen receptors. Nat. Chem. Biol. 14, 317–324 (2018).

12. Liang, W.-C. et al. Cross-species Vascular Endothelial Growth Factor (VEGF)-blocking Antibodies Completely Inhibit the Growth of Human Tumor Xenografts and Measure the Contribution of Stromal VEGF *. J. Biol. Chem. 281, 951–961 (2006).

13. Neuzillet, C. et al. Targeting the TGFβ pathway for cancer therapy. Pharmacol. Ther. 147, 22–31 (2015).

14. Metelli, A. et al. Surface Expression of TGFβ Docking Receptor GARP Promotes Oncogenesis and Immune Tolerance in Breast Cancer. Cancer Res. 76, 7106–7117 (2016).

15. Sahtoe, D. D. et al. Reconfigurable asymmetric protein assemblies through implicit negative design. Science 375, eabj7662 (2022).

16. Edman, N. I. et al. Modulation of FGF pathway signaling and vascular differentiation using designed oligomeric assemblies. 2023.03.14.532666 Preprint at 10.1101/2023.03.14.532666 (2023).

17. Gordon, W. R., Arnett, K. L. & Blacklow, S. C. The molecular logic of Notch signaling – a structural and biochemical perspective. J. Cell Sci. 121, 3109–3119 (2008).

18. Toda, S. et al. Engineering synthetic morphogen systems that can program multicellular patterning. Science 370, 327–331 (2020).

19. Steinbuck, M. P., Arakcheeva, K. & Winandy, S. Novel TCR-Mediated Mechanisms of Notch Activation and Signaling. J. Immunol. 200, 997–1007 (2018).

20. Maesako, M., Houser, M. C. Q., Turchyna, Y., Wolfe, M. S. & Berezovska, O. Presenilin/γ-Secretase Activity Is Located in Acidic Compartments of Live Neurons. J. Neurosci. 42, 145–154 (2022).

21. Zhao, L., Zhao, J., Zhong, K., Tong, A. & Jia, D. Targeted protein degradation: mechanisms, strategies and application. Signal Transduct. Target. Ther. 7, 1–13 (2022).

22. Zhao, C. et al. Receptor Heterodimerization Modulates Endocytosis through Collaborative and Competitive Mechanisms. Biophys. J. 117, 646–658 (2019).

23. Rollins, C. T. et al. A ligand-reversible dimerization system for controlling protein–protein interactions. Proc. Natl. Acad. Sci. U. S. A. 97, 7096–7101 (2000).

24. Salter, A. I. et al. Comparative analysis of TCR and CAR signaling informs CAR designs with superior antigen sensitivity and in vivo function. Sci. Signal. 14, eabe2606 (2021).

25. Hammill, J. A. et al. Designed ankyrin repeat proteins are effective targeting elements for chimeric antigen receptors. J. Immunother. Cancer 3, 55 (2015).

26. Hammill, J. A. et al. A Cross-Reactive Small Protein Binding Domain Provides a Model to Study Off-Tumor CAR-T Cell Toxicity. Mol. Ther. - Oncolytics 17, 278–292 (2020).

27. Morgan, R. A. et al. Case Report of a Serious Adverse Event Following the Administration of T Cells Transduced With a Chimeric Antigen Receptor Recognizing ERBB2. Mol. Ther. 18, 843–851 (2010).

28. Prantner, A. M. et al. Anti-Mesothelin Nanobodies for Both Conventional and Nanoparticle-Based Biomedical Applications. J. Biomed. Nanotechnol. 11, 1201–1212 (2015).

29. Williams, J. Z. et al. Precise T cell recognition programs designed by transcriptionally linking multiple receptors. Science 370, 1099–1104 (2020).

